# Misspellings or “miscellings”-non-verifiable cell lines in cancer research publications

**DOI:** 10.1101/2024.02.29.582220

**Authors:** Danielle J. Oste, Pranujan Pathmendra, Reese A. K. Richardson, Gracen Johnson, Yida Ao, Maya D. Arya, Naomi R. Enochs, Muhammed Hussein, Jinghan Kang, Aaron Lee, Jonathan J. Danon, Guillaume Cabanac, Cyril Labbé, Amanda Capes Davis, Thomas Stoeger, Jennifer A. Byrne

**Affiliations:** School of Medical Sciences, Faculty of Medicine and Health, The University of Sydney, NSW, Australia; Sydney School of Veterinary Science, Faculty of Science, The University of Sydney, NSW, Australia; Department of Chemical and Biological Engineering, Northwestern University, Evanston, IL, USA; School of Chemistry, Faculty of Science, The University of Sydney, NSW, Australia; IRIT UMR 5505 CNRS, University of Toulouse, Toulouse, France; Institut Universitaire de France (IUF), Paris, France; Université Grenoble Alpes, CNRS, Grenoble INP, Laboratoire d’Informatique de Grenoble, Grenoble, France; CellBank Australia, Children’s Medical Research Institute, The University of Sydney, Westmead, NSW, Australia; Feinberg School of Medicine in the Division of Pulmonary and Critical Care Medicine, Northwestern University, Chicago, IL, USA; The Potocsnak Longevity Institute, Northwestern University, Chicago, IL, USA; Simpson Querrey Lung Institute for Translational Science, Chicago, IL, USA; NSW Health Statewide Biobank, NSW Health Pathology, Camperdown, NSW, Australia

## Abstract

Reproducible laboratory research relies on correctly identified reagents. We have previously described human gene research papers with wrongly identified nucleotide sequence reagent(s), including papers studying *miR-145*. Manually verifying reagent identities in more recent *miR-145* papers found 20/36 (56%) and 6/36 (17%) *miR-145* papers with misidentified nucleotide sequence reagent(s) and human cell line(s), respectively. We also found 5 cell line identifiers in two *miR-145* papers with wrongly identified nucleotide sequences and cell lines, and 18 identifiers published elsewhere that did not correspond to indexed cell lines. These cell line identifiers were described as non-verifiable, as their identities appeared uncertain. Studying 420 papers that mentioned 8 different non-verifiable cell line identifier(s) found 235 papers (56%) that appeared to refer to BGC-803, BSG-803, BSG-823, GSE-1, HGC-7901, HGC-803 and/or MGC-823 as independent cell lines. We could not find publications describing how these cell lines were established, and they were not indexed in claimed externally accessible cell line repositories. While some papers stated that STR profiles had been generated for BGC-803, GSE-1 and/or MGC-823 cells, no STR profiles were identified. In summary, non-verifiable human cell lines represent new challenges to research reproducibility and require further investigation to clarify their identities.

**Novelty and Impact Statement:** Through verifying reagent identities in research publications, our team found 23 non-verifiable human cell line identifiers, most of which could represent misspellings of contaminated cancer cell lines. Of 8 identifiers studied in detail, 7 non-verifiable identifiers were unexpectedly referred to as independent cell lines across 235 publications. We therefore describe a process “miscelling”, where published cell lines lack descriptions of how they were established, cannot be found in claimed external repositories and lack STR profiles.

## Introduction

Preclinical research aims to produce reliable observations that can be further tested or extended through translational studies. A variety of resources can serve as reagents in biomedical experiments, including cell lines, antibodies to detect proteins of interest, and oligonucleotide reagents to analyze specific genomic sequences or transcripts (1).

While it is commonly assumed that antibodies, cell lines and nucleotide sequence reagents are correctly identified, many studies now indicate that experimental reagents can be wrongly identified in publications. In addition to problems of antibody cross-reactivity and variable performance (2), ELISA kits have been found to include antibodies that detect different targets from those claimed (3, 4). Independent studies have described research papers where the identities of some nucleotide sequence reagents do not match their descriptions (5–17), including oligonucleotide probes on microarrays (18, 19), and shRNAs in libraries (20). Finally, due to cross-contamination and/or uncertainty over donor origins, many human cell lines are known to be wrongly identified (21–25). These wrongly identified reagents can produce a range of downstream consequences, including failed attempts to reproduce published results (3, 5, 7, 11, 16), and potentially incorrect interpretations of preclinical data leading to misdirected translational research (10, 12, 25).

The capacity to verify reagent identities can help researchers to assess whether published results have been correctly reported. Verifiable reagents include short oligonucleotides and peptides, as their sequences are typically disclosed (10, 13). Because oligonucleotide and peptide sequences usually cannot be identified by eye, they need to be linked with gene or protein identifiers (13, 15, 17). Reagent identities can therefore be verified by querying nucleotide or amino acid sequences against DNA/RNA or protein sequence databases, and then comparing predicted and claimed reagent identities (10, 13–15, 17). Similarly, cell lines are named using identifiers composed of short strings of letters and/or numbers (22, 26). These identifiers can be used to query knowledgebases such as Cellosaurus (26) to check whether (i) the claimed identity matches the indexed identity, and (ii) cell lines are known or suspected to be cross-contaminated or otherwise misclassified. Cell line knowledgebases can also help researchers to navigate ambiguities of cell line nomenclature, such as identifiers that do not reflect the cell line’s biological origin, variations in syntax and punctuation, and identical or very similar identifiers being used to denote different cell lines (22, 26–28). In addition to matching RRIDs to cell line names (28), Cellosaurus also indexes recognized synonyms for cell line identifiers, and in some cases, potentially misspelled identifiers noted in the literature (26).

This project sought to extend an earlier study that examined the proportions of human gene research publications that described wrongly identified nucleotide sequences (15). Park et al. (15) screened >11,700 human gene research papers with the semi-automated tool Seek & Blastn that verifies the identities of nucleotide sequence reagents that are claimed to target human transcripts and genomic sequences (13). Papers screened by Seek & Blastn were distributed across 5 literature corpora, including a corpus of original papers that studied human *miR-145* in preclinical cancer models (15). Screening 163 *miR-145* papers with Seek & Blastn identified 31 (19%) *miR-145* papers with one or more wrongly identified nucleotide sequences (15). Other human gene research papers that studied different miRs were also found to include wrongly identified nucleotide sequence reagents (15). In addition to wrongly identified nucleotide sequences (15, 17), other issues have been highlighted in the preclinical miR literature, such as experiments where targeted miRs might not be expressed at physiologically relevant levels (29–31). For example, although *miR-145* has been repeatedly studied in human cancer cell lines of epithelial origin, *miR-145* is not expressed in epithelial cells and would not be expected to serve as a critical regulator in many human cancer cell lines (32, 33).

To determine whether *miR-145* papers with wrongly identified nucleotide sequences have been published since the analyses of Park et al. (15), the present study manually verified the identities of nucleotide sequence reagents (10, 17) in *miR-145* papers from 2020-2022. We also examined the human cell lines described in recent *miR-145* papers, to determine whether any experiments were conducted with cross-contaminated or misclassified cell lines. These approaches found numerous *miR-145* papers with wrongly identified nucleotide sequences and human cell lines. Furthermore, in two *miR-145* papers with incorrect nucleotide sequences and cell lines, we also identified 5 cell line identifiers that did not correspond to indexed cell lines. Two identifiers (BGC-803, BSG-823) were recognized misspellings of the contaminated human cell lines MGC-803 and BGC-823, respectively (26). The remaining identifiers (BSG-823, GSE-1, TIE-3) were not indexed by Cellosaurus, but could also have represented misspelled versions of contaminated cell lines (26). We collectively referred to these identifiers as being non-verifiable (NV), as their identities were unclear. While all NV identifiers were expected to represent misspellings of similarly named cell lines, the BGC-803, BSG-823 and GSE-1 identifiers appeared to be described as independent cell lines in one *miR-145* paper. As 5 NV identifiers were found in just two *miR-145* papers, we undertook further analyses to find other NV cell line identifiers and understand how a subset of these identifiers have been described in the literature.

## Materials and Methods

### Identification of miR-145 corpus

To identify recent *miR-145* papers not studied by Park et al. (15), Web of Science was searched using the following parameters: Title=“miR-145” AND Topic=“Cancer” AND All Fields=“Human” AND Document Type=“Article” for articles published from 01 January 2020 to 01 September 2022. Search results were exported into Google Sheets, articles were visually screened to ensure that each paper had studied human *miR-145*, and the title, journal, publisher, publication year, PubMed ID (PMID) and journal impact factor (JIF) (34) for the publication year were recorded. Human cancer type(s) studied were identified by visual inspection of text. Country of origin was determined according to the affiliations of at least half of the authors (15, 17). Publications were judged to be hospital-affiliated (15, 17) or university-affiliated if at least half of authors listed relevant affiliations. In the case of no majority, the first author’s affiliation(s) determined affiliation status (17).

### Verification of nucleotide sequence reagent identities

Nucleotide sequence identities were verified if reagents (i) claimed to target a non-modified human gene, transcript, or genomic sequence, or to represent a non-targeting control and (ii) were listed in the article and/or supplementary text or tables. Identities of nucleotide sequence reagents were not verified if they were (i) claimed to target gene transcripts or genomic sequences in species other than human, (ii) claimed to target human genes that could not be identified in either GeneCards (35) or miRBase (36), or (iii) shown in formats such that sequences could not be directly extracted, for example within image files.

Nucleotide sequence identities were verified as described (14, 15), except for claimed circRNA targeting reagents, which were verified as described by Pathmendra et al. (17). Nucleotide sequence identities were verified by individual co-authors and re-checked by DJO, whereas all nucleotide sequences verified by DJO were re-checked by other co-authors. DJO also re-checked (i) a randomly-selected subset (>10%) of nucleotide sequences and (ii) all wrongly identified nucleotide sequences, in consultation with PP and/or JAB.

Nucleotide sequence reagents that were indicated to be wrongly identified were classified into one of 3 error types (13–15, 17): (i) reagents claimed to target a specific human gene, transcript or genomic sequence, but indicated to be non-targeting in human; (ii) targeting reagents predicted to target a different human gene, transcript or genomic sequence from that claimed (including circRNA targeting reagents predicted to not discriminate circular from linear transcripts); and (iii) claimed non-targeting reagents predicted to target a human gene transcript.

### Verification of cell line identities

Cell line identifiers in *miR-145* papers were extracted using copy/paste functions and used to query Cellosaurus (26). Cell lines indexed by Cellosaurus as being (potentially) contaminated with a different cell line or otherwise misclassified were flagged if the article did not recognize (i) the cell line’s contaminated/misclassified status or (ii) use of a contaminated/misclassified cell line as a study limitation.

### Literature searches employing cell line identifiers

Cell line identifiers were employed as keywords to search Google Scholar (Figure S1). Keywords were employed with and without a dash (“-”) between alphabetic and numeric identifier components, where “-” was also recognized as a space by Google Scholar. Where search results and GeneCards (35) indicated that any cell line identifier corresponded to a human gene (27, 37), identifiers were combined with the terms “cells” and “cell line”, ie “TIE-3 cells”, “TIE-3 cell line”. Where Cellosaurus did not index an identifier as a human cell line, or only recognized an identifier as a misspelled human cell line identifier (26), the identifier was referred to as non-verifiable (NV). Publications retrieved by NV cell line identifiers (+/- “cells” and “cell line”) were prioritized for analysis according to their best match ranking. Where NV cell line identifiers retrieved up to 50 sources, all publications were selected for further analysis. Otherwise, 50-200 publications were selected from the first 6-20 pages of Google Scholar results, with a focus on top-ranked results.

Publications were further analyzed if they could be downloaded in full or sourced through University of Sydney library services and were written in English. Text search functions were used to identify the queried cell line identifier, and all figures and tables were visually inspected. If publications included the queried identifier, the following information was extracted: (i) article type (ii) PMID, DOI or other identifier, (iii) publication year, (iv) journal, (v) publisher, (vi) JIF for the publication year (if available), (vii) publication title, (viii) country affiliation, (ix) hospital/ university affiliation, (x) details of any post-publication notices (corrections, expression of concerns, retractions), (xi) whether the queried identifier was used in experiments and/or referenced, (xii) the claimed cell type (eg. gastric cancer), (xiii) whether the queried identifier was listed with other cell lines, and their identities, (xiv) cell line sources, if provided and (xv) whether cell line identities were checked using short tandem repeat (STR) profiling.

Google Scholar results were also triaged by date, to identify the earliest publications that had referred to queried identifiers. Publications were subjected to further analyses if they could be sourced as described above, and if at least the abstract was written in English and mentioned the queried cell line identifier. Publications were visually inspected for descriptions of cell line establishment (1, 23) and other publication features were recorded as described above.

### Literature analyses of NV and similar human cell line identifiers

Google Scholar searches were conducted with NV human cell line identifiers paired with known human cell line identifiers, where the NV identifier was either a recognized misspelling of a human cell line identifier (26) and/or was similar to and/or occurred in association with a similarly named human cell line. One index paper listed two NV identifiers (BGC-803, BSG-823) with similarly named cell lines (HS-746T, BSG-823, MKN-28, 9811, BGC-803, MGC-803, BGC-823). This identifier cluster was also used as a search term (Data file S1). Publications were triaged and analysed as described above. Where cell line sources represented repositories with accessible online catalogues, catalogues were queried with NV cell line identifiers written (i) as single words and (ii) with a space or “-” between the alphabetic and numeric components. For each queried catalogue, a human cell line identifier (eg. MCF-7) was employed as a positive control.

As some publications that referred to NV cell line identifiers stated that cell line identities had been confirmed using STR profiling, additional Google Scholar searches combined NV cell line identifiers with the terms “STR” and “short tandem repeat”. All publications that referred to NV cell line identifiers were searched for references to STR profiling and STR profiling results.

As some *miR-145* papers referred to NV cell line identifiers as independent cell lines (see Results), publications that referred to NV and similarly named human cell line identifiers were inspected to determine whether NV cell line identifiers were (i) likely to represent misspelled versions of human cell line identifiers, or (ii) referred to as independent cell lines. A NV cell line identifier was indicated to represent a *misspelling* if the NV identifier (i) was used alternately with similarly named human cell line(s) such that identifiers were never directly connected, either in the same sentence (eg. “and”) or adjacent sentences (eg. “also”, “in contrast”); (ii) did not appear in any list of cell lines with any similarly named cell line; and (iii) were not included in any single experiment with any similarly named cell line. In contrast, NV cell line identifiers were indicated to represent *independent cell lines* if (i) the NV identifier was used in the absence of similarly-named human cell line(s); (ii) NV and similarly-named human cell line identifiers were included in any list of cell lines studied in the paper; (iii) results for NV and similarly-named human cell line(s) were shown in the same experiment(s); and/or (iv) NV and similarly-named human cell line identifiers were directly connected, either in the same or adjacent sentences.

### Representation of cell line identifiers

We have chosen to show all cell line identifiers in the text with a dash (-) between alphabetical and numeric components, and to list identifiers in alphabetical order, for improved readability.

### Citation analyses

Google Scholar citations were collected on 29 January, 2024.

## Results

### miR-145 corpus and nucleotide sequence analyses

We identified 36 original publications that studied human *miR-145*, referred to human cancer and were published between 01 January 2020 and 01 September 2022. Most (24/36, 67%) *miR-145* papers described nucleotide sequence reagents, where these 24 *miR-145* articles were published in 21 journals and examined *miR-145* in the context of 15 human cancer types (Table S1).

The 24 *miR-145* papers described 339 nucleotide sequence reagents, of which almost one quarter (83/339, 24%) were predicted to be wrongly identified (Table 1). Most incorrect nucleotide sequences were claimed targeting reagents that were either predicted to be non-targeting in human (43/83, 52%), or to target a different human gene or genomic sequence from that claimed (39/83, 47%) (Table S2, Data file S2). Wrongly identified nucleotide sequences were found in over half (20/36, 56%) of *miR-145* papers, and in most (20/24, 83%) *miR-145* papers that described nucleotide sequence reagents, with a median of 3 (range 1-16) wrongly identified sequences/ paper (Table 1). The 20 *miR-145* papers with wrongly identified nucleotide sequence(s) have been cited 382 times (Table S1).

**Table 1.** Verification of reagent identities in human *miR-145* papers published from 2020-2022.

|  | **Nucleotide sequence reagents** | **Cell lines<sup>a</sup>** | **Total** |
| --- | --- | --- | --- |
| Number of *miR-145* papers with reagents whose identities could be verified | n=24 | n=21 | n=24 |
| Number of reagents per *miR-145* paper, median (range) | 13 (2-54) | 5 (1-13) | 18 (4-56) |
| Number of *miR-145* papers according to country of origin | China (n=21), Brazil (n=1), Taiwan (n=1), Thailand (n=1) | China (n=19), Taiwan (n=1), Thailand (n=1) | China (n=21), Brazil (n=1), Taiwan (n=1), Thailand (n=1) |
| Proportion (percentage) of wrongly identified reagents | 83/339 (24%) | 15/107 (14%) | 98/446 (22%) |
| Proportion (percentage) of *miR-145* papers with wrongly identified reagents | 20/24 (83%) | 6/21 (29%) | 20/24 (83%) |
| Number of wrongly identified reagents per paper, median (range) | 3 (1-16) | 3 (1-4) | 5 (1-16) |
| Number of papers with wrongly identified reagents according to country of origin | China (n=19), Thailand (n=1) | China (n=6) | China (n=19), Thailand (n=1) |
| Proportion (percentage) of papers with wrongly identified reagents that were affiliated with hospitals | 16/20 (80%) | 5/6 (83%) | 16/20 (80%) |

<sup>a</sup> Refers to cell lines indexed by Cellosaurus (release 47, October 2023), does not include non-verifiable cell line identifiers

### Cross-contaminated and misclassified cell lines

A total of 21 *miR-145* papers described experiments in cell lines, where all papers also described nucleotide sequence reagent(s) (Table S1). These 21 *miR-145* papers described 91 cell lines, with a median of 5 (range 1-13) cell lines/paper (Table 1, Data file S2). Fourteen percent (15/107) of cell lines were wrongly identified (Table 1), and almost one third (6/21, 29%) of *miR-145* papers that described cell lines included at least one wrongly identified cell line. All wrongly identified cell lines were claimed human cancer cell lines, where most have been found to be contaminated by HeLa and/or other cancer cell lines (38–41) (Table S3).

### Identification of non-verifiable (NV) cell line identifiers

Two *miR-145* papers that described wrongly identified nucleotide sequence(s) and cell line(s) also described cell line identifier(s) that were either recognized misspellings of the contaminated cell lines MGC-203 and BGC-823 (BGC-803, BSG-823) (26), and/or not indexed by Cellosaurus (BSG-803, GSE-1, TIE-3) (Tables S4, S5). We commonly referred to recognized misspelled identifiers and non-indexed identifiers (26) as non-verifiable (NV), as their identities appeared uncertain.

As 5 NV identifiers were found in two *miR-145* papers, we reasoned that other NV cell line identifiers might exist in the literature. Google Scholar searches successively identified 23 different NV cell line identifiers (Tables S4 and S5), many of which were similar to contaminated cell lines such as BGC-823, MGC-803 or SGC-7901 (Table S5). We analysed 8 NV identifiers in detail, including the 5 NV cell line identifiers found in *miR-145* papers (Tables S4 and S5). In the papers where NV identifiers were first identified (referred to henceforth as index papers), 4 NV identifiers (BGC-803, BSG-823, GSE-1, HGC-7901) were referred to as independent cell lines, whereas the remaining 4 NV identifiers appeared to represent misspellings of known cell lines (Tables S4 and S5). Furthermore, 3 NV identifiers (BGC-803, BSG-823, GSE-1) were claimed to have been obtained from a cell line repository with an online catalogue (Table S4) yet querying this repository catalogue did not identify these 3 cell line identifiers.

### Descriptions of NV identifiers in the research literature

To understand how the 8 NV cell line identifiers have been described in the literature, we assessed 420 articles that referred to at least one of these NV identifiers (Data file S1). In 185/420 (44%) papers, NV identifier(s) appeared to represent misspellings of similarly named cell lines, which were typically contaminated cancer cell lines (38–41). For example, in the 79 papers that referred to both GSE-1 and GES-1, GSE-1 appeared to consistently represent a misspelling of GES-1 (Data file S1). Similarly, in the single paper that referred to TIE-3 as a cell line identifier, TIE-3 appeared to be a misspelling of TE-3 (Table S4).

While many instances of NV identifiers seemed to represent misspellings, more than half (235/420, 56%) of papers appeared to describe at least one of 7 NV identifiers (BGC-803, BSG-803, BSG-823, GSE-1, HGC-7901, HGC-803, and/or MGC-823) as independent cell lines (Tables 2 and 3, Data file S1). Some original papers that described the results of experiments involving NV cell line identifier(s) referred to NV identifier(s) without reference to any similarly named human cell line (26). For example, 34 original papers referred to GSE-1 as a cell line identifier, without any mention of the GES-1 cell line (Table 2). In other original papers, NV and similarly named cell lines were (i) included in lists of cell lines employed in experiments (for example: “The human gastric cancer cell lines … MGC-823, MGC-803…were used in this study”), (ii) used in the same experiments according to information provided in the text, and/or (iii) shown together as results for the same experiment(s) in figures and/or tables (Data file S1). The numbers of original papers that described NV identifiers as independent cell lines ranged from one paper (BSG-803) to 116 papers (BGC-803). Papers that did not describe experiments with NV cell lines, including 36 literature reviews, also referred to NV identifiers as independent cell lines (Table 3). In these papers, NV identifiers were found in the absence of a similarly named cell line or were paired with similar cell line identifiers, either in literature review text, or in introduction or discussion sections of original papers (Data file S1). Papers describing NV identifiers as independent cell lines were published between 2004-2023 in fields including cancer research, biochemistry, chemistry and drug discovery, and in journals with impact factors ranging from 0.2-16.8 (Tables 2 and 3).

**Table 2.** Original papers that describe experiments where non-verifiable (NV) cell line identifier(s) were described as independent cell line(s)

| **NV cell line identifier** | **Original papers referring to NV identifier as independent cell line<sup>a</sup>** | **Publication years range** | **Number of individual journals/publishers** | **Journal Impact Factor range** | **Most frequent country of origin proportion (%)** | **Most frequent institution type proportion (%)** | **Cell line repository sources** | **Papers referring to derivation of STR profiles proportion (%)** |
| --- | --- | --- | --- | --- | --- | --- | --- | --- |
| BGC-803 | n=116 | 2006-2023 | n=82/ n=34 | 0.2-12.7 | China 115/116 (99%) | Hospital 73/116 (63%) | Named collaborator/ institute/ laboratory; ATCC; BeNa Culture Collection (Beijing, China); Cell Bank of the Chinese Academy of Sciences; Type Culture Collection of Chinese Academy of Sciences | 4/116 (3%) |
| BSG-803 | n=1 | 2020 | n=1/ n=1 | 4.1 | China 1/1 (100%) | Hospital 1/1 (100%) | Not stated | 0/1 (0%) |
| BSG-823 | n=14 | 2015-2022 | n=13/ n=11 | 2.5-6.4 | China 14/14 (100%) | Hospital 12/14 (86%) | Named collaborating institute; Cell Bank of the Chinese Academy of Sciences; National Infrastructure of Cell Line Resources of China | 0/14 (0%) |
| GSE-1 | n=34 | 2004-2023 | n=30/ n=16 | 2.0-9.7 | China 33/34 (97%) | Hospital 27/34 (79%) | Named collaborating institute; ATCC; BeNa Culture Collection; China Center for Type Culture Collection | 2/34 (6%) |
| HGC-803 | n=3 | 2016-2018 | n=3/ n=3 | 1.7-4.3 | China 3/3 (100%) | Hospital 3/3 (100%) | ATCC; Cell Bank of the Chinese Academy of Sciences | 0/3 (0%) |
| HGC-7901 | n=7 | 2015-2023 | n=7/ n=5 | 2.1-8.5 | China 7/7 (100%) | Hospital 4/7 (57%) | Cell Bank of Chinese Academy of Sciences | 0/7 (0%) |
| MGC-823 | n=34 | 2005-2023 | n=30/ n=16 | 0.3-11.2 | China 33/34 (97%) | Hospital 23/34 (68%) | Named collaborator or collaborating institute; ATCC; Cell Bank of Chinese Academy of Sciences; China Center for Type Culture Collection | 5/34 (15%) |

<sup>a</sup>Original papers that referred to experiments that employed NV cell line(s) (i) without reference to any similarly-named cell line and/or (ii) with a similarly-named cell line, such that NV cell line(s) were referred to as independent cell line(s)

**Table 3.** Literature reviews, commentaries, book chapters, original articles and preprints where non-verifiable (NV) cell line identifier(s) were referred to as independent cell line(s) in the absence of experimental results.

| **NV cell line identifier** | **Number of papers (literature reviews) referring to NV identifier as independent cell line** | **Publication years range** | **Number of individual journals/publishers** | **Journal Impact Factor range** | **Most frequent country of origin proportion (%)** | **Most frequent institution type proportion (%)** |
| --- | --- | --- | --- | --- | --- | --- |
| BGC-803 | n=27 (n=22) | 2009-2023 | n=22/ n=8 | 1.8-13.6 | China<br>11/21 (41%) | University<br>22/27 (81%) |
| BSG-823 | n=8 (n=7) | 2016-2023 | n=7/ n=5 | 2.8-13.0 | China<br>4/8 (50%) | University<br>5/8 (63%) |
| GSE-1 | n=2 (n=2) | 2023 | n=2/ n=2 | No<br>information | N/A<sup>a</sup> | University<br>2/2 (100%) |
| HGC-7901 | n=1 (n=1) | 2020 | n=1/ n=1 | 6.2 | China 1/1 (100%) | Hospital 1/1 (100%) |
| HGC-803 | n=4 (n=2) | 2019-2023 | n=3/ n=3 | 1.1-6.6 | China<br>2/4 (50%) | N/A<sup>a</sup> |
| MGC-823 | n=2 (n=2) | 2019-2021 | n=2/ n=2 | 7.8-16.8 | N/A<sup>b</sup> | University<br>2/2 (100%) |

<sup>a</sup>N/A in relation to country/ institution indicates no majority

### NV cell line establishment, sources and STR profiling results

To clarify the origins of NV cell line identifiers, we analyzed the earliest publications that mentioned NV identifiers and/or referred to NV identifiers as cell lines. The earliest papers that mentioned NV cell line identifiers were published between 1988-2020, whereas the first papers that referred to NV identifiers as independent cell lines were published between 2004-2020 (Figure 1). Whereas the BSG-803, BSG-823, GSE-1 and HGC-7901 identifiers appeared to be first mentioned as independent cell lines, BGC-803, HGC-803, and MGC-823 were referred to as independent cell lines 1-17 years after they were first mentioned in the literature (Figure 1). No earliest paper that we could find described how the 7 NV cell lines were established or referenced other cell line establishment papers.

**Figure 1.**
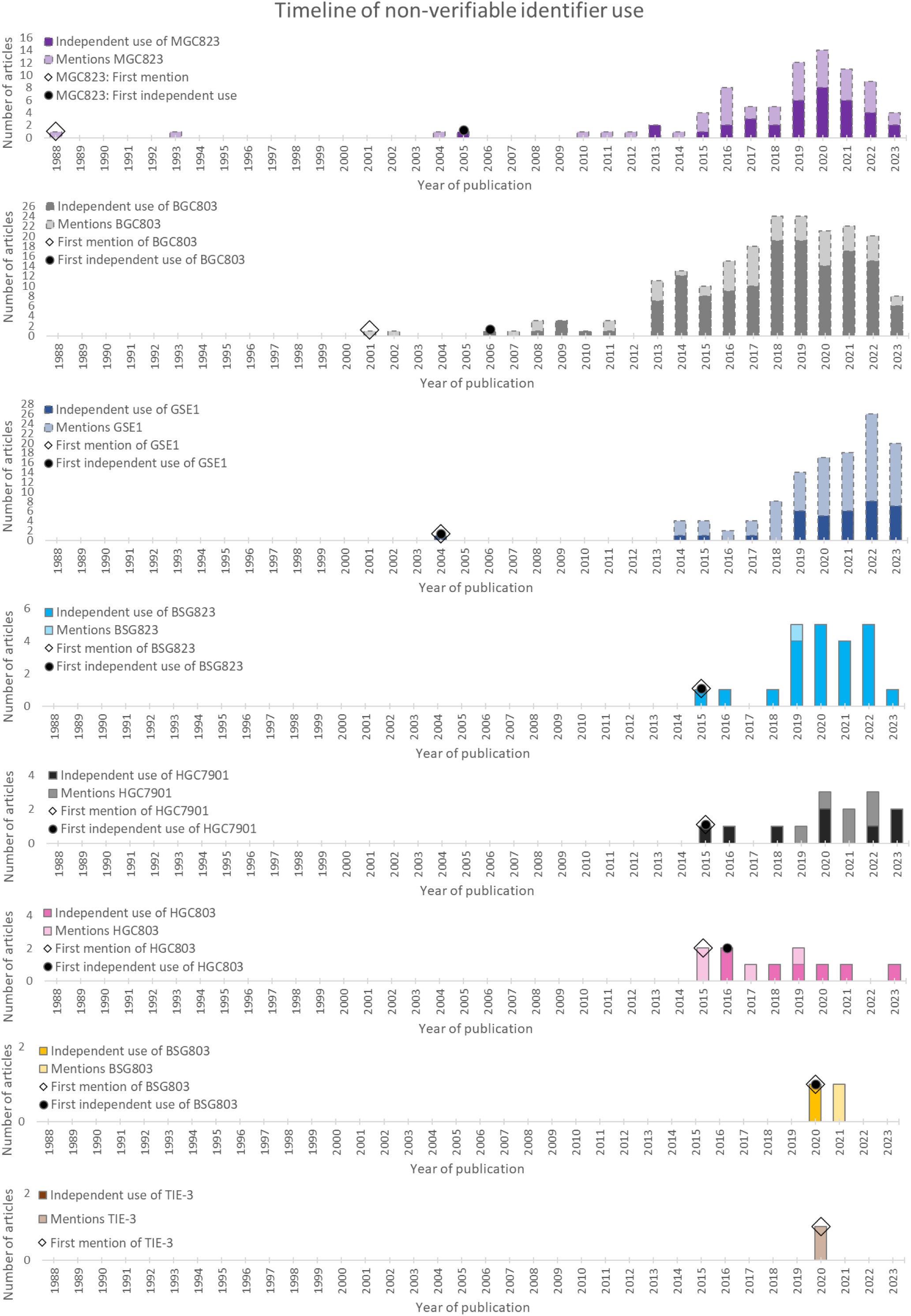
Bar graphs showing numbers of papers (Y axes) that refer to the indicated NV cell line identifiers (shown at top left) per year (X axis). Light shaded bars indicate papers that mention NV cell line identifiers as likely misspellings, whereas darker bars indicate papers in which NV cell line identifiers were referred to as independent cell lines. Broken lines indicate publication numbers that derived from studying a subset of papers (BGC-803, GSE-1, MGC-823), as described in the Methods. Open diamonds indicate the earliest publication that mentioned each identifier, whereas filled circles indicate the earliest paper to refer to a NV identifier as an independent cell line. Graphs are ranked from top according to year of earliest publication (1988-2020).

As we did not find 3 NV identifiers in the described cell line repository in one index paper (Table S4), we extracted all cell line sources described in the 420 papers (Data file S1). In papers where NV identifiers were referred to as independent cell lines, 4 NV cell lines (BGC-803, GSE-1, HGC-803, MGC-823) were described as sourced from the American Type Culture Collection (ATCC) (42). Six NV cell lines (BGC-803, BSG-823, GSE-1, HGC-7901, HGC-803, MGC-823) were described as sourced from the Chinese National Infrastructure of Cell Line Resource (43), and 3 NV cell lines (BGC-803, GSE-1, MGC-823) were described as sourced from the China Center for Type Culture Collection (38–41). Searching the online catalogues of these 3 repositories did not identify the claimed NV identifiers or cell lines (Table S5).

A small minority (16/420, 4%) of papers that mentioned NV cell line identifiers described the use of STR profiling to confirm cell line identities (Table 2, Data file S1). These 16 papers either described STR profiling to confirm the identities of named NV cell lines or included overarching statements that were presumed to refer to all cell lines. Although 11/16 papers referred to 3 NV cell line identifiers (BGC-803, GSE-1 and/or MGC-823) as independent cell lines, no paper either showed or described STR profiling results for any NV cell line (Data file S1).

## Discussion

This study commenced by checking nucleotide sequence reagent identities in a cohort of human *miR-145* papers. Having previously identified wrongly identified nucleotide sequences in 19% human *miR-145* papers using Seek & Blastn screening (15), manual screening revealed at least one wrongly identified nucleotide sequence in over half (56%) *miR-145* papers published between 2020-2022, and in 83% *miR-145* papers that described nucleotide sequences (Table 1). While these higher proportions likely reflect the use of manual screening (17), these results demonstrate that papers with wrongly identified nucleotide sequences represent an ongoing challenge to the *miR-145* field (15).

Just under one third of relevant *miR-145* papers also described wrongly identified cell line(s) (Table 1), in addition to wrongly identified nucleotide sequences (Table S1). Unexpectedly, two *miR-145* papers that described both wrongly identified nucleotide sequence(s) and cell line(s) also included 5 NV cell line identifiers (Table S4). Two NV identifiers (BGC-803, BSG-823) were recognized misspellings (26) of the contaminated cell lines BGC-823 or MGC-803 (38–41), whereas the BSG-803, GSE-1 and TIE-3 identifiers were similar to the contaminated cell lines BGC-823/ MGC-803 (38–41), GES-1 (39, 41) and TE-3, respectively (44). While all NV identifiers could therefore represent cell line identifier misspellings, we were surprised to find that BGC-803, BSG-823 and GSE-1 were referred to as independent cell lines. As 5 NV cell line identifiers were found in two *miR-145* papers, and paralleled previous descriptions of NV circular RNAs (17, 45), we sought to understand whether other NV cell line identifiers could be found in the literature and how a subset of these NV identifiers have been described.

Literature searches found 18 other NV cell line identifiers in addition to those found in *miR-145* papers (Table S5). The frequency of these NV identifiers varied, where some were found only once as cell line identifiers, whereas others were found in hundreds of Google Scholar sources. Studying 8 NV cell line identifiers across 420 papers showed that all 8 NV identifiers could represent misspellings of indexed cell lines, which were typically HeLa-contaminated cancer cell lines. However, 56% papers referred to a total of 7 different NV identifiers as independent cell lines. In some cases, the NV identifier was used in the absence of any similarly named cell line, whereas other papers mentioned NV identifiers in association with similarly named cell lines, such that NV identifiers appeared to refer to independent cell lines. These NV cell line identifiers have been mentioned in the literature as early as 1988 and have been referred to as independent cell lines since 2004 (Figure 1). Nonetheless, we could not find any papers that described how these NV cell lines were established. Six different NV cell lines were stated to have been sourced from named cell line repositories, including ATCC, yet checking online catalogues did not identify these NV cell lines. We also could not identify any published STR profiles for BGC-803, GSE-1 and MGC-823 cells, despite papers stating that these STR profiles had been confirmed. These NV cell lines were also not described by studies describing STR profiles for similarly named contaminated cell lines (38–41).

### Limitations

We recognize that this study has several limitations. Although we attempted to study all papers that mentioned the identifiers BSG-803, BSG-823, HGC-803, HGC-7901 and TIE-3, we may have still missed some relevant publications. For example, as GSE-1 and TIE-3 also represent human genes, our use of additional search terms to restrict results may have excluded relevant papers. We conducted literature searches for NV cell line identifiers during 2023, and so we may have missed some relevant papers published late in 2023 (Figure 1). We also recognize that as we only screened papers with at least some English text, we could have failed to identify papers that described how NV cell lines were established, and/or the results of STR profiling. Due to numbers of available publications, we did not analyze all papers that mentioned the BGC-803, GSE-1 and MGC-823 identifiers. We did not study all NV identifiers that we found (Table S5) and we made no attempt to find all NV cell line identifiers that might exist in the literature.

While claimed NV cell lines could not be found in 3 external repositories with online catalogues, including ATCC (42), we recognize that stocks of these and other NV cell lines might be held elsewhere, including repositories lacking online catalogues. We also recognize that at least some papers that referred to NV identifiers without reference to similarly named cell line(s) could have employed the NV identifier as a misspelling, and that similar explanations could apply to some other descriptions of independent NV cell lines. Nonetheless, given that many NV cell line identifiers could represent misspellings of cross-contaminated cancer cell lines, descriptions of any intended cell lines could remain problematic.

### Significance and next steps

While we recognize that some NV identifiers represent misspellings of similarly named cell lines, our results indicate that some misspelled identifiers can also gain new identities as independent cell lines. This seems to occur without published descriptions of how cell lines were established, STR profiles, or making cell lines available through external suppliers, a process that we refer to as “miscelling” (Figure 2). While some NV cell lines and their corresponding STR profiles may simply have never been described in the literature, our inability to locate claimed cell lines in external repositories raises doubts as to their status. As 5 NV identifiers were also found in *miR-145* publications with wrongly identified nucleotide sequence(s), NV cell lines could represent further indications that at least some experiments were not performed as described.

**Figure 2.**
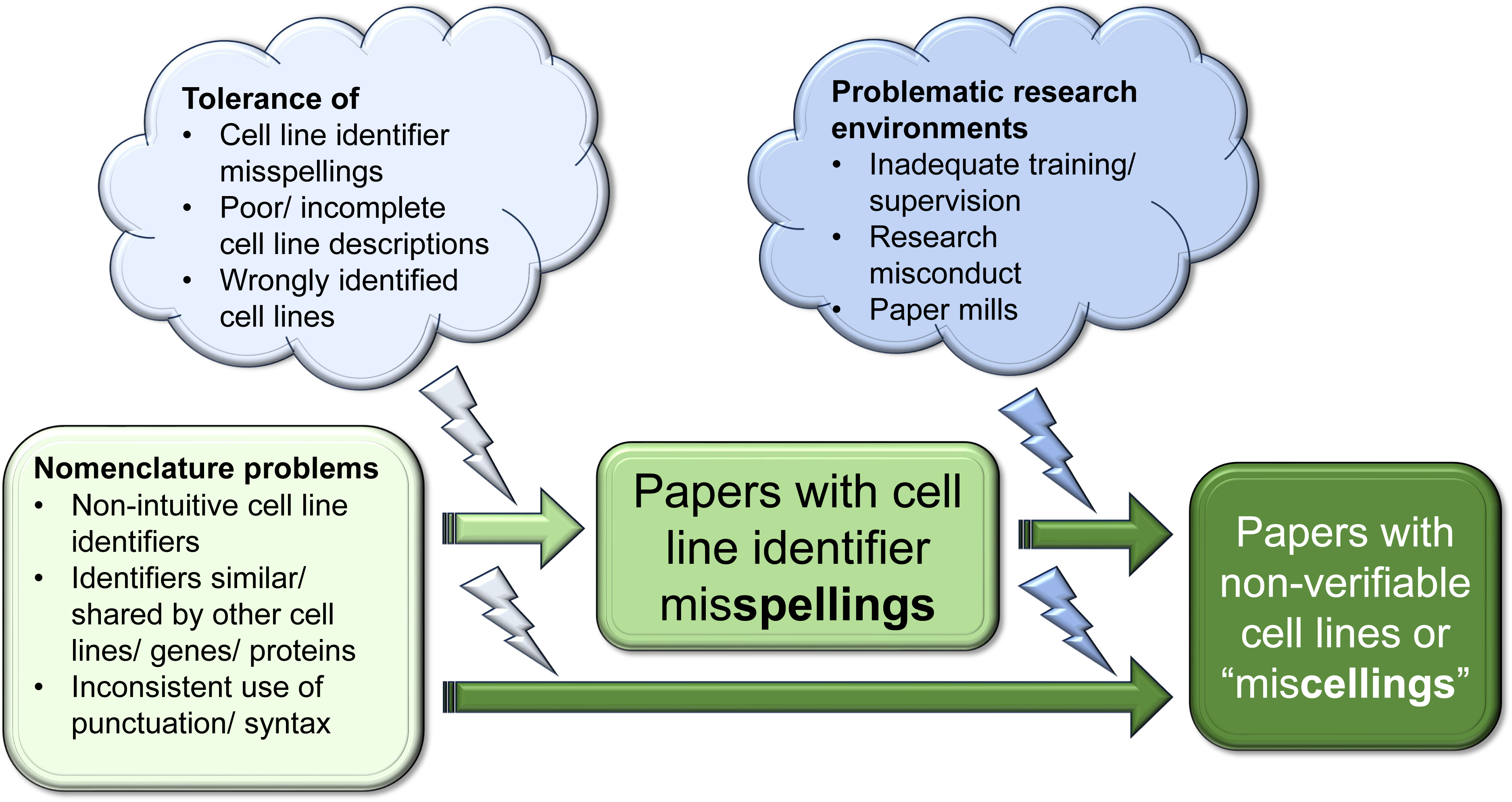
Summary of factors that could predispose to cell line identifier misspellings and NV cell lines or “miscellings”, recognizing that papers that appear to refer to NV cell lines can either (i) temporally follow publications where NV identifiers are likely to represent simple misspellings, or (ii) appear de novo.

We recommend that NV cell line identities be clarified as soon as possible, either by disclosing existing STR profiles and/or supplying cell line stocks to independent groups for STR profiling and phenotypic testing. While we could not identify sources for NV cell lines, teams that have described these cell lines could provide samples for testing, where dates of cell line stocks should predate published experiments. Testing cell line stocks from different sources would have the added advantage of allowing STR profiles for multiple cell line stocks to be directly compared (38–41).

Repeated references to misspelled cell line identifiers in Cellosaurus (26), combined with the many NV identifiers found in this study, suggest that other NV cell line identifiers remain to be discovered. Misspelled and other NV cell line identifiers may be difficult to notice through reading, particularly where cell line identifiers are non-intuitive and can be written with different punctation and/or syntax (22, 26–28) (Figure 2). Furthermore, as misspelled versions of familiar commercial logos can be recognized as if they were correct (46), even expert readers might not notice subtle identifier misspellings. Most NV identifiers described by this study were found by undergraduate students working on short-term publication integrity projects. Similar projects could be conducted in future, particularly as publication integrity projects can be scaled according to available time and resources and are suitable as individual or group projects.

It will also be important to understand the conditions that allow NV cell lines to arise and become more frequent in the literature (Figure 2). For example, frequent cell line misspellings in publications could lead researchers to accept and use misspelled identifiers, particularly in settings with limited cell culture knowledge and experience (Figure 2). We suggest steps to reduce descriptions of NV cell lines (Table 4), where some have also been proposed to reduce descriptions of contaminated or misidentified cell lines (23, 26, 28, 47, 48, 50). Requiring authors to include the RRID (28) for each cell line within the Materials and Methods section would improve the quality of published cell line identifiers and could also reduce descriptions of NV cell lines. To raise awareness of NV cell lines, we suggest creating a dynamic registry of NV and misspelled cell lines. Adding misspelled, NV and wrongly identified cell lines to screening tools (49) could improve their detection in manuscripts and publications. We recommend a zero-tolerance approach for manuscripts that describe misspelled and NV cell line identifiers, where such manuscripts should not be sent for peer review. It has been recommended that publications that describe experiments with misidentified cell lines should be flagged with editorial notes or expressions of concern (23, 50). This should extend to all publications that describe NV cell lines. If the identities and origins of cell line(s) cannot be verified, papers that have described experiments with NV cell lines should be considered for retraction.

**Table 4.** Actions to prevent descriptions of non-verifiable (NV) cell lines in research publications

| **Actions to reduce descriptions of NV cell lines** | **Previously described in context of wrongly identified cell lines<sup>a</sup>** | **Stakeholders** |
| --- | --- | --- |
| Establish dynamic register of misspelled cell line identifiers and NV cell lines |  | Researchers, research funding organizations |
| Create screening tools/ plug-ins to detected misspelled, NV and wrongly identified cell lines | 28, 49 |  |
| Insist on use of correct cell line identifiers in research publications. Zero tolerance for misspelled cell line identifiers. | 22, 47, 48 | Authors, journals, publishers, peer reviewers |
| Cell lines to be described in easily searchable publication sections, eg title and/or abstract | 23 |  |
| Cell lines to be described by at least two identifiers, eg cell line identifier + RRID | 24, 28, 48 |  |
| All source(s) of published cell lines to be fully disclosed | 23 |  |
| All cell lines described in results to also be clearly described in the methods |  |  |
| Publications describing new cell line to include description of donor origin, method of establishment, culture conditions, phenotyping and genotyping data including STR profile | 1, 23 |  |
| Verification of all cell line identities prior to peer review | 23, 24 | Journals, publishers |
| Immediate publication of expressions of concern or editorial notes for papers that describe non-verifiable cell lines | 23, 50 |  |

<sup>a</sup>References that have described or proposed similar approach(es) to reduce descriptions of cross-contaminated or misidentified cell lines

### Summary

As reported for other identifiers that lack intrinsic visual sense (51), cell line identifiers may be prone to being wrongly written or misspelled (22, 26). Although some NV identifiers are likely to represent misspellings of similarly named cell lines, our results indicate that some misspelled cell line identifiers can gain new identities as independent cell lines, a process that we describe as “miscelling”. As NV cell lines represent challenges to research reproducibility, further research is needed to confirm the identities of NV cell lines and determine whether other NV cell lines have been described. In the meantime, NV cell lines call for urgent improvements to published descriptions of human cell lines, to ensure the transparency of cell line models employed in biomedical research.

## Acknowledgements

JAB, ACD and CL gratefully acknowledge funding from the National Health and Medical Research Council of Australia (NHMRC) Ideas grant ID APP1184263. PP is supported by a Research Training Program scholarship at the University of Sydney. RAKR gratefully acknowledges support from the Dr. John N. Nicholson fellowship from Northwestern University and Moderna Inc. “Identifying bias and improving reproducibility in RNA-seq computational pipelines”. JJD is supported by an NHMRC Investigator Grant (GNT2008066). TS is supported by NIH grant AG068544.

## Author contributions

**Conceptualization**: JAB, GC; CL; **Methodology:** JAB, DJO, PP, ACD; GC, CL; **Formal analysis:** DJO, PP, RAKR, GJ, YA, MDA, NRE, MH, JK, AL, JAB; **Writing - original draft preparation:** JAB, DJO; **Writing - review and editing:** DJO, PP, RAKR, GJ, YA, MDA, NRE, MH, JK, AL, JD, GC, CL, ACD, JAB; **Funding acquisition:** JAB, ACD, CL; **Supervision:** JAB, JD, TS.

## Declarations

**Ethical Approval:** Not applicable

## Data Availability Statement

All data generated or analysed during this study are included in the manuscript and its Supplementary Information files. All information extracted from or about analysed publications, as well as Google Scholar citation data, is available within the public domain.

## List of Supplementary Figures/ Tables/ Data files

**Data file S1** Non-verifiable cell line identifier analysis

**Data file S2** *miR-145* cohort reagent analysis

**Figure S1.**
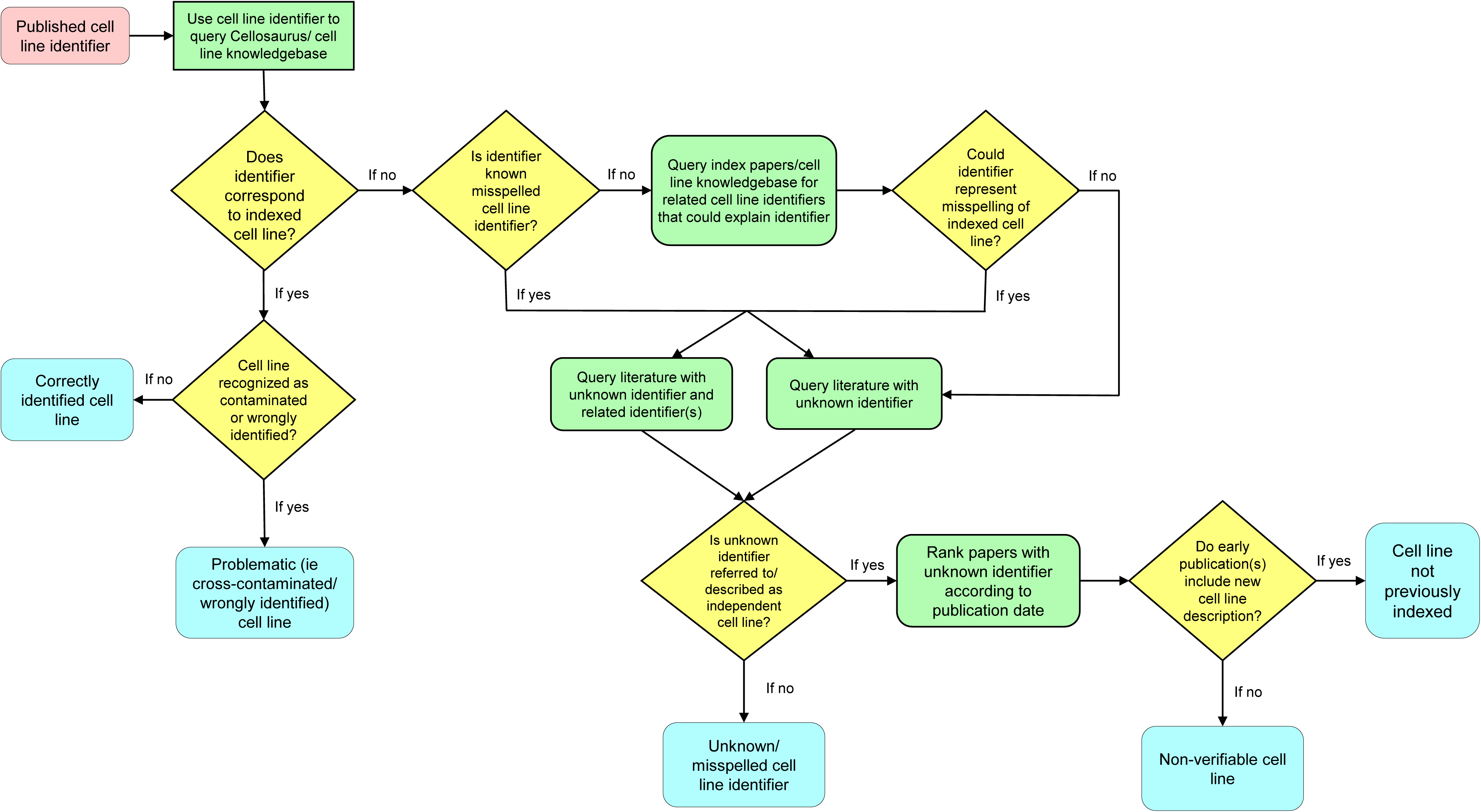
Flow chart summarising workflows used to manually verify cell line identities

**Table S1.**
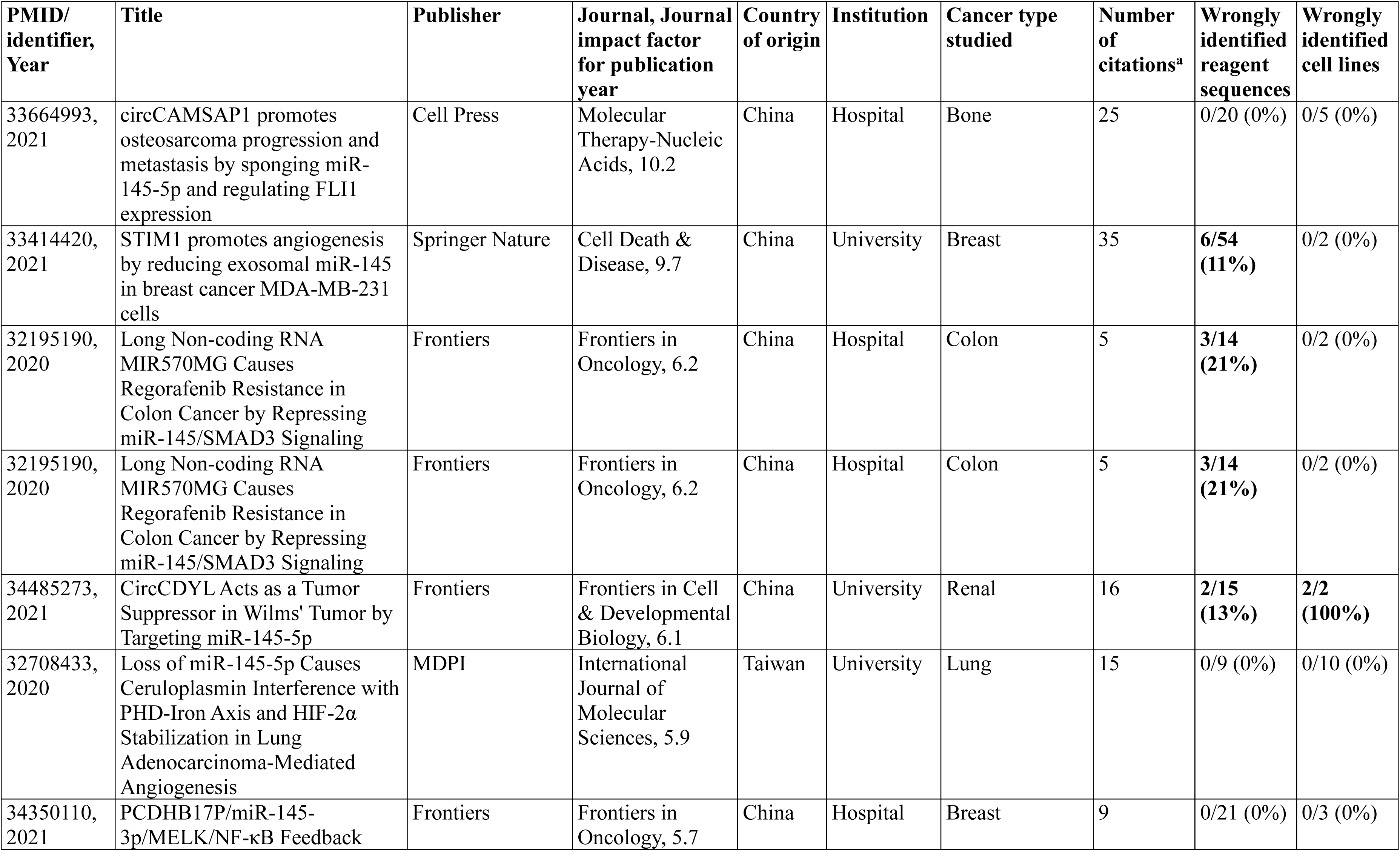
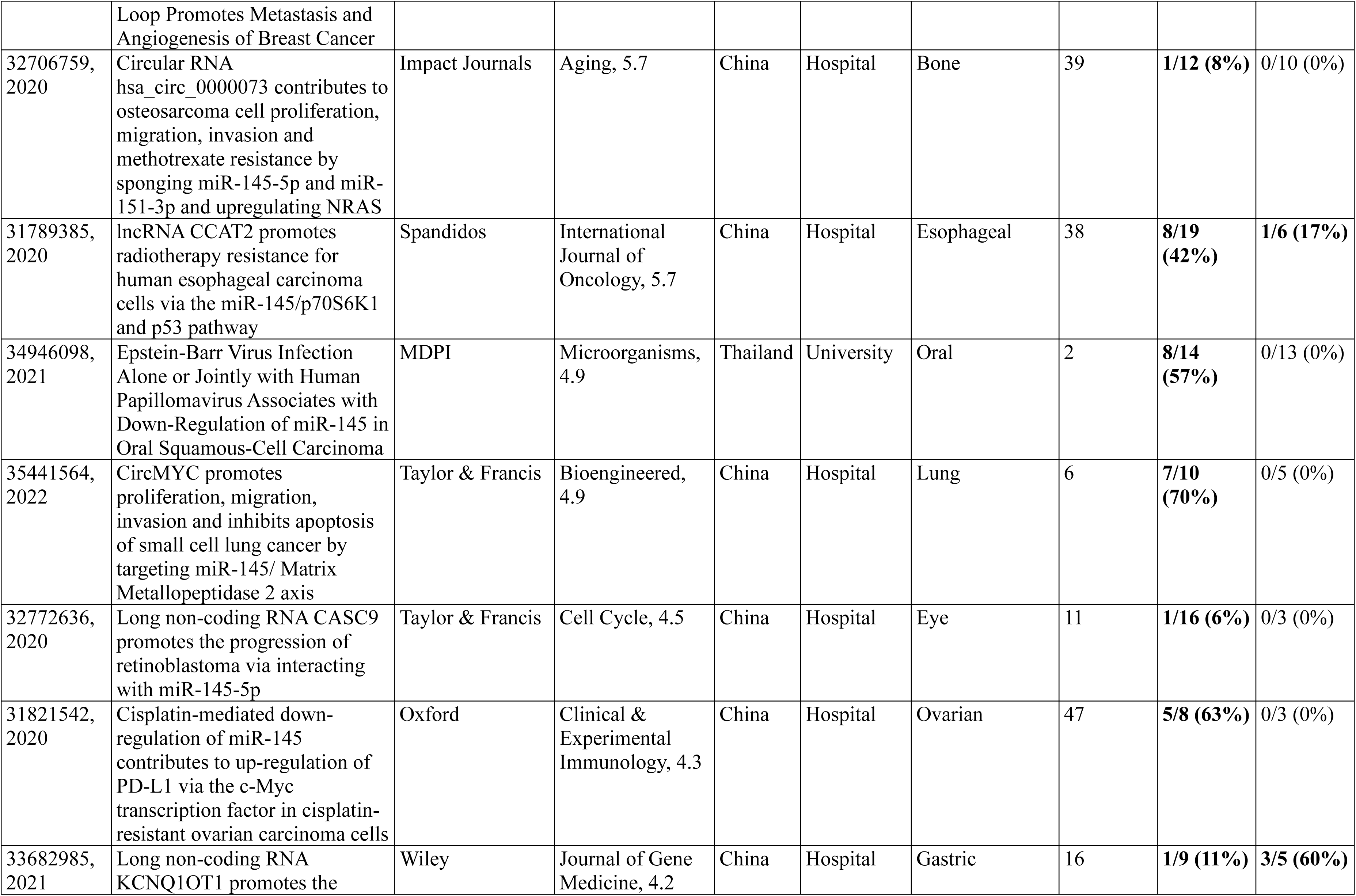
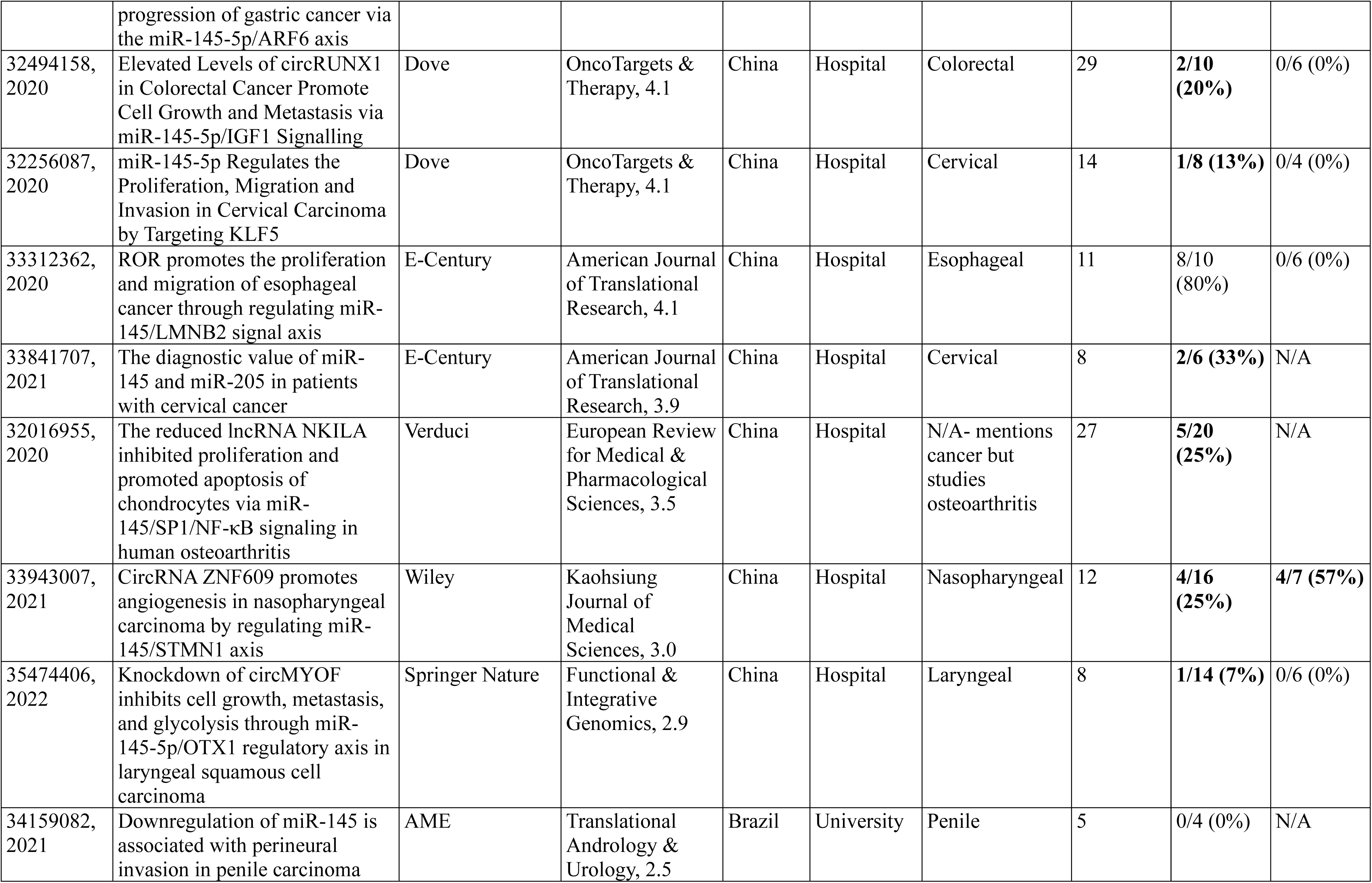
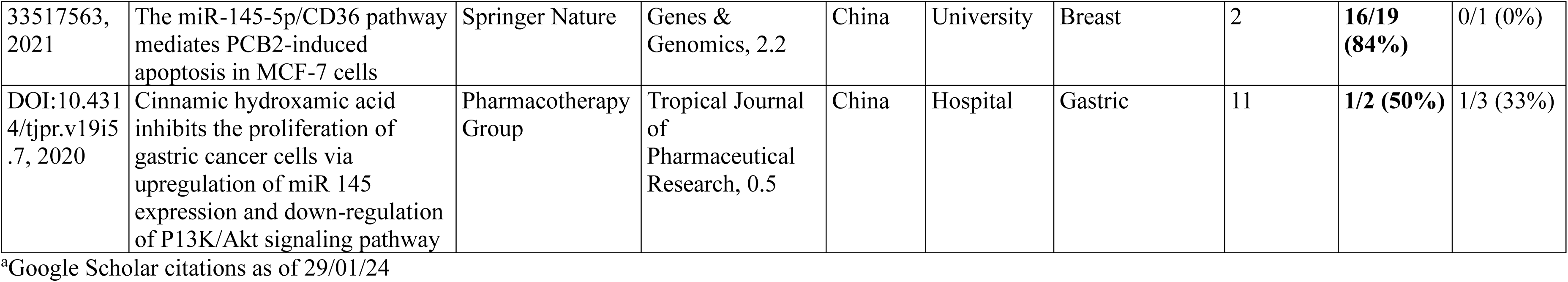
*miR-145* papers that described nucleotide sequence reagents, ranked according to journal impact factor.

**Table S2.** Wrongly identified reagents in *miR-145* papers published from 2020-2022.

| **Reagent type** | **Proportion (percentage)** |
| --- | --- |
| **Wrongly identified nucleotide sequence reagents (n=20 *miR-145* papers)** | 83/339 (24%) |
| Claimed targeting reagents predicted to be non-targeting in human | 43/83 (52%) |
| Claimed targeting reagents predicted to target a different human gene or genomic sequences | 39/83 (47%) |
| Claimed non-targeting reagents predicted to target a human gene | 1/83 (1%) |
| Non-verifiable nucleotide sequence reagents | 8 |
| **Problematic cell lines (n=6 *miR-145* papers)** | 15/107 (14%) |
| Cross-contaminated cell lines | 12/15 (80%) |
| Misclassified cell lines | 3/15 (20%) |
| **Non-verifiable cell line identifiers** | 5 |

**Table S3.** Wrongly identified cell lines in *miR-145* papers published from 2020-2022, ranked by PMID and then DOI.

| PMID | Claimed identity | Cell line identifier | Verification results | Verified identity from Cellosaurus<sup>a</sup> |
| --- | --- | --- | --- | --- |
| 31789385 | Esophageal cancer | TE-3 | Contaminated | Identical to TE-2, TE-7, TE-12 and TE-13 |
| 33682985 | Human gastric cancer | BGC823 | Contaminated | HeLa derivative |
|  | Human gastric cancer | MKN28 | Contaminated | MKN74 derivative |
|  | Human gastric cancer | MGC803 | Possibly contaminated | HeLa and cell of unknown origin hybrid |
| 33858418 | Human hepatocellular carcinoma | Bel-7402 | Contaminated | HeLa derivative |
|  | Human normal hepatic cell line | L02 | Contaminated | HeLa derivative |
|  | Human hepatocellular carcinoma | SMMC-7721 | Contaminated | HeLa derivative |
|  | Human hepatocellular carcinoma | HepG2 | Misclassified | Hepatoblastoma |
| 33943007 | Nasopharyngeal carcinoma | 5-8F | Contaminated | HeLa and cell of unknown origin hybrid |
|  | Nasopharyngeal carcinoma | CNE1 | Contaminated | HeLa and cell of unknown origin hybrid; Identical to CNE-2 |
|  | Nasopharyngeal carcinoma | HONE-1 | Possibly contaminated | HeLa derivative and cell of unknown origin |
|  | Nasopharyngeal carcinoma | HNE1 | Possibly contaminated | High similarity to HeLa and HNE-2 |
| 34485273 | Wilms' tumor (nephroblastoma) | SK-NEP-1 | Misclassified | Ewing Sarcoma |
|  | Wilms' tumor (nephroblastoma) | G401 | Misclassified | Kidney rhabdoid tumor |
| DOI:10.4314/tjpr.v19i5.7 | Human gastric cancer | SGC-7901 | Contaminated | HeLa derivative |

<sup>a</sup> Cellosaurus release 47 (October 2023)

**Table S4.** Earliest/ index publications that describe 8 non-verifiable (NV) cell line identifiers, ranked by NV identifier alphabetical order.

| Earliest/<br>Index (Cell<br>line<br>Identifier(s)) | PubMed ID,<br>Publication<br>year | Journal<br>(Journal<br>Impact Factor)<br>Publisher | Publication title | NV cell line<br>identifier(s) -<br>described as<br>independent cell<br>line(s)? Yes/No | Source of NV cell<br>line | Wrongly<br>identified<br>cell line(s) | Wrongly<br>identified<br>nucleotide<br>sequences |
| --- | --- | --- | --- | --- | --- | --- | --- |
| Index<br>(BGC803;<br>BSG803,<br>BSG823,<br>GSE1) | 33682985,<br>2021 | Journal of Gene<br>Medicine (4.2)<br>Wiley | Long non-coding RNA<br>KCNQ1OT1 promotes the<br>progression of gastric cancer via<br>the miR-145-5p/ARF6 axis | • BGC803-Yes<br>• BSG803-No<br>• BSG823-Yes<br>• GSE1-Yes | • BGC803,<br>BSG823, GSE1:<br>Cell Bank of the<br>Chinese Academy<br>of Sciences | • BGC823<br>• MGC803<br>• MKN28 | 1/9 (11%) |
| Earliest/<br>Index<br>(HGC7901) | 25043310,<br>2015 | Oncogene (7.9)<br>Springer Nature | MicroRNA-25 promotes gastric<br>cancer migration, invasion and<br>proliferation by directly targeting<br>transducer of ERBB2, 1 and<br>correlates with poor survival | • HGC7901-Yes | • HGC7901: source<br>not provided | • BGC-823<br>• SGC-7901 | 0/11 (0%) |
| Earliest/<br>Index<br>(HGC803) | 26102366,<br>2015<br>(Retracted<br>2023) | PLoS One (3.1)<br>PLoS | MicroRNA-137 Contributes to<br>Dampened Tumorigenesis in<br>Human Gastric Cancer by<br>Targeting AKT2 | • HGC803-No | N/A<sup>a</sup> | • MGC803<br>• SGC7901 | 0/12 (0%) |
| Earliest/<br>Index<br>(MGC823) | https://doi.org/10.1007/BF02997636,<br>1988 | Chinese Journal<br>of Cancer<br>Research (N/A)<br>Chinese Journal<br>Cancer<br>Research Co | Production of antitumor human<br>monoclonal antibodies by human-<br>human or mouse-human<br>hybridoma | • 475 A- Yes<br>• Clue- No<br>• E 109- Yes<br>• Keto- No<br>• MGC823- No<br>• Mnk-45- No<br>• MNK-75- No<br>• Natasch- Yes | • 475 A, E 109:<br>named individuals<br>• Natasch: source<br>not provided | • 7402<br>• BC-1<br>• BGC823<br>• M-21<br>• MGC803<br>• SGC7901 | Nucleotide<br>sequences<br>not<br>included |
| Earliest/<br>Index (TIE-3) | 31789385,<br>2020 | International<br>Journal of<br>Oncology (5.7)<br>Spandidos | lncRNA CCAT2 promotes<br>radiotherapy resistance for human<br>esophageal carcinoma cells via<br>the miR-145/p70S6K1 and p53<br>pathway | • TIE-3-No | N/A<sup>a</sup> | • TE-3 | 8/19 (42%) |

<sup>a</sup>Not described as independent cell line

**Table S5.** Non-verifiable (NV) human cell line identifiers found in the present study, ranked by alphabetical order.

| NV cell line identifier<sup>a</sup> | Assumed/<br>claimed<br>identity | Recognised<br>as misspelled<br>identifier by<br>CelloSaurus? | Similarly named cell<br>line- identity<br>Letters/ numbers in bold<br>differ in NV identifier | Index paper<br>PMID/ DOI | Google Scholar results<br>n= (search string)<br>January-February 2024 | Found in ATCC/<br>cellbank.org.cn/<br>CCTCC?<sup>c</sup> (Y/N) |
| --- | --- | --- | --- | --- | --- | --- |
| 475 A | Breast cancer | No | Unclear<sup>b</sup> | doi.org/10.1007/BF02997636 | 0 (“475 A cell line”) | N/ N/ N |
| BGC-230 | Gastric cancer | No | BGC-**823**- contaminated | PMID: 30416662 | 1 (“BGC230 cells”) | N/ N/ N |
| BGC-7901 | Gastric cancer | No | SGC-7901- contaminated | PMID: 26671739 | 48 (BGC7901 cells) | N/ N/ N |
| **BGC-803** | Gastric cancer | **Yes** | BGC-**823**- contaminated | PMID: 33682985 | 505 (BGC803 cells) | N/ N/ N |
| BGC-829 | Gastric cancer | No | BGC-**823**- contaminated | PMID: 8381684 | 4 (BGC829 cells) | N/ N/ N |
| **BSG-803** | Gastric cancer | No | BGC-**823**- contaminated | PMID: 33682985 | 2 (BSG803 cells) | N/ N/ N |
| **BSG-823** | Gastric cancer | **Yes** | BGC-**823**- contaminated |  | 34 (BSG823 cells) | N/ N/ N |
| Clue | Lung cancer | No | Unclear<sup>b</sup> | doi.org/10.1007/BF02997636 | 0 (“Clue cell line”) | N/ N/ N |
| E 109 | Esophageal<br>cancer | No | Unclear<sup>b</sup> |  | 0 (“E 109 cell line”) | N/ N/ N |
| **GSE1** | Gastric<br>epithelial | No | GES-1- contaminated | PMID: 33682985 | 13 (“GSE1 cells”) | N/ N/ N |
| **HGC-7901** | Gastric cancer | No | SGC-7901- contaminated | PMID: 25043310 | 16 (HGC7901 cells) | N/ N/ N |
| **HGC-803** | Gastric cancer | No | **MGC**-803-? contaminated | PMID: 26102366 | 34 (HGC803 cells) | N/ N/ N |
| HGC-823 | Gastric cancer | No | **BGC**-823- contaminated | PMID: 26817615 | 1 (HGC823 cells) | N/ N/ N |
| Keto | Gastric cancer | No | Unclear<sup>b</sup> | doi.org/10.1007/BF02997636 | 2 (“Keto cell line”) | N/ N/ N |
| **MGC823** | Gastric cancer | No | MGC-**803**-? contaminated | https://doi.org/10.1007/BF02997636 | 158 (MGC823 cells) | N/ N/ N |
| MKN-75 | Gastric cancer | No | MKN-**45**- gastric cancer | PMID: 12782760 | 27 (“MKN75” cells) | N/ N/ N |
| Mnk-45 | Gastric cancer | **Yes** | MKN-45- gastric cancer | doi.org/10.1007/BF02997636 | 609 (“Mnk45” cells) | N/ N/ N |
| MNK-75 | Gastric cancer | No | MKN-**45**- gastric cancer |  | 3 (“MNK-75” cells) | N/ N/ N |
| Natasch | Neuroglioma | No | Unclear<sup>b</sup> |  | 10 (“Natasch” cell line) | N/ N/ N |
| SGC-27 | Gastric cancer | No | **HGC**-27- gastric cancer | PMID: 28404932 | 103 (“SGC-27” cells) | N/ N/ N |
| SGC-803 | Gastric cancer | No | **MGC**-803-? contaminated | PMID: 33349155 | 35 (SGC-803 cells) | N/ N/ N |
| SGC-823 | Gastric cancer | No | **BGC**-823- contaminated | PMID: 32864003 | 39 (SGC-823 cells) | N/ N/ N |
| **TIE-3** | Esophageal<br>cancer | No | TE-3- contaminated | PMID: 31789385 | 3 (“TIE-3 cells”) | N/ N/ N |

<sup>a</sup>NV cell line identifiers shown in bold were analysed in depth (Tables 2, 3, Table S4)
<sup>b</sup>No similarly named cell line was found
<sup>c</sup>All repository catalogue searches used syntax variations (+/- space/hyphen) of NV cell line identifiers, where relevant

